# Using dual EEG to analyse event-locked changes in child-adult neural connectivity

**DOI:** 10.1101/2021.06.15.448573

**Authors:** I. Marriott Haresign, E. Phillips, M. Whitehorn, L. Goupil, S.V. Wass

## Abstract

Current approaches typically measure the connectivity between interacting physiological systems as a time-invariant property. This approach obscures crucial information about how connectivity between interacting systems is established and maintained. Here, we describe methods, and present computational algorithms, that will allow researchers to address this deficit. We focus on how two different approaches to measuring connectivity, namely concurrent (e.g., power correlations, phase locking) and sequential (e.g., Granger causality), can be applied to three aspects of the brain signal, namely amplitude, power, and phase. We guide the reader through worked examples using mainly simulated data on how to leverage these methods to measure changes in interbrain connectivity between adults and children/infants relative to events identified within continuous EEG data during a free-flowing naturalistic interaction. For each, we aim to provide a detailed explanation of the interpretation of the analysis and how they can be usefully used when studying early social interactions.

## Section 1 Introduction

Behavioural evidence points to clear social influences on how infants pay attention (Yu & Smith, 2016) and learn (Kuhl et al., 2003) during early social engagement. But we currently understand little about how these interpersonal influences are supported in the brain (Wass et al., 2020; Redcay & Schilbach, 2019; Redcay & Warnell, 2018; Hoehl et al., 2021). Hyper scanning is a method of simultaneously acquiring neural activity from two or more individuals that allows insights into this question (Dumas et al., 2010; Schilbach et al, 2013). Hyper scanning approaches are often paralleled with an emphasis on using more free-flowing ’naturalistic’ study designs that record brain activity during real-life interactions – rather than studying neural responses to repetitive and un-ecological trial-based structured tasks administered via a computer screen.

Recently, research with non-human animals (e.g., Kingsbury et al 2019; Zhang & Yartsev, 2019) and human adults (Liu et al., 2018; Redcay & Schilbach, 2019), as well as research with children/infants using fNIRS (Nguyen et al., 2021; Piazza et al., 2021; Reindl et al., 2018) has started to use hyperscanning to uncover complex patterns of interbrain connectivity (IBC) that may develop during social interaction. More recently, EEG research with child/infant populations has also started to address this. From this research, we know that bidirectional Granger-causal influences between infants’ and adults’ neural activity are greater in theta (3-6Hz) and alpha frequency bands (6-9Hz) during mutual gaze than during moments of non-mutual/ averted gaze (Leong et al., 2017). We also know that both undirected (phase locking) and directed (spectral Granger-causal) IBC patterns in the theta and alpha bands are higher when adults model positive emotions during social interaction than when adults model negative emotions (Santamaria et al., 2020). These findings suggest that consistent with fNIRS studies that have shown IBC patterns over longer temporal scales (e.g., Piazza et al., 2021; Nguyen, et al., 2020; 2021), IBC may also be discernible at a more fine-grained, sub-second scale studied using EEG.

All these approaches used thus far, however, share one fundamental limitation. Hyper-scanning researchers typically calculate the amount of IBC observed between two interacting physiological systems averaged across whole conditions (Perez et al., 2017; Leong et al., 2017) and even whole interactions (e.g., Kinreich et al., 2017) and then compare IBC values between different conditions (e.g. averaging all mutual gaze moments and comparing them with non-mutual gaze) (Leong et al., 2017), or they correlate findings with an outcome variable (e.g., learning) (Leong et al., 2019).

Effectively, therefore, these approaches produce an index of IBC that includes information on how connectivity varies by frequency (e.g. Leong et al., 2017) and by scalp topography (e.g. Santamaria et al., 2020) – but which excludes information on how IBC fluctuates over time. This omission is, we argue, fundamentally hinders our understanding of how real-life social interactions are substantiated in the brain. The same observation largely holds for most of the research in adult populations, where similar points have been raised concerning the limitations of current approaches (e.g., Novembre & Ianetti, 2021; Moreau & Dumas, 2021).

### 1.1 The importance of the missing time dimension

Studies using event-related potentials (ERPs) have shown that even young infants’ brains show millisecond-level sensitivity to ostensive cues (e.g. Farroni et al., 2002; Hoehl & Striano, 2008; 2010, Quadrelli et al., 2019). But this research is all unidirectional: it examines how the recipient of an ostensive signal is influenced by the ‘sender’ of the signal. Very little research has examined the fine-grained temporal dynamics of early social interaction from a bidirectional perspective: by examining how ostensive cues affect the inter-relationship between both partners’ brain activity (Wass et al., 2020).

One of the few early studies that has measured this found that, in the 3-9Hz range, neural activity in one partner consistently predicts the other partner’s neural activity more strongly during direct compared with indirect gaze (Leong et al., 2017). But how is it mechanistically possible for two brains influence each other over such fine-grained temporal scales?

I. First, it is possible that certain behavioural events in social interaction could lead to transient changes in spectral power (e.g., Grossman et al., 2007) and/or phase (e.g., Makeig et al., 2004) in both the ‘sender’ and the ‘receiver’ of the social cue, leading to increases in IBC (see e.g. section 2.2.1, 2.2.2). This mechanism would be similar to those documented for neural entrainment to speech (e.g., Doelling et al., 2014). These behavioural events might be mutual gaze onsets, or vocalisations (e.g., Lachat, 2012; Müller & Lindenberger, 2019). It is also possible that the magnitude of these changes could be mediated by factors including attention (e.g., Golumbic et al., 2013), comprehension (e.g., Pérez et al., 2019), and environmental factors such partner familiarity (e.g., Reindl et al., 2021, in press). According to this model, IBC would peak around these salient events and decrease thereafter.
II. Second, it is possible that response preparation or anticipation (e.g., Hamilton, 2021; Hirsch et al., 2017; Kirkland, 2020) and mutual prediction (Hamilton 2021) might lead to concurrent transient changes in either power or phase in both partners (e.g., Mandel et al., 2016; Bögels, 2019), causing increases in IBC (see e.g. section 2.2.3, 2.2.4). It is also possible that the magnitude of these changes could be affected by factors such as the amount (e.g., Nguyen, et al, 2021) and quality (e.g., Bloom, 1988) of turn-taking. According to this hypothesis, IBC would peak around these ‘handover’ moments and decrease before and after.
III. Third, it is possible that continuous intra brain changes, that are not locked to behavioural events, could lead to gradual changes in IBC. This might be substantiated in three ways. First, it might take the form of direct ‘neural mimicry’. For example, Kingsbury and colleagues (2019) used *in vivo* electrophysiological recordings to show populations of cells in the dorsomedial PFC that show similar activity when performing an action as when watching it be performed by someone else (Kingsbury et al., 2019). Second, it might be driven by shared entrainment to aspects of the shared environment, such as language or body cues. For example, Simony and colleagues showed inter-participant entrainment was increased when participants had a shared understanding of a story (Simony et al., 2016). Third, they might be driven by mutual prediction even in the absence of explicit turn-taking (Hamilton, 2021; Kingsbury et al., 2019). Again, these changes might take the form of changes in power: increases in spectral power throughout an event (e.g., a look/ episode of attention) can increase signal-to-noise ratios and cause increases in sequential IBC (as we show in section 3.2.1). This is conceivable as, for example, infant theta power increases through an attentional episode (e.g. Jones et al., 2020). Alternatively, gradual changes in phase, such as the adjustment of the peak frequency of neural oscillations, could lead to increases in concurrent IBC (section 3.2.2). This is conceivable as, for example, peak alpha frequency can be modulated by task demands (e.g., Samaha & Postle, 2015; Wutz et al., 2018), and recent accounts have theorised that cross-spectrum frequency adjustment at stimulus onset might be a mechanism behind how ERPs are generated (e.g., Burgess 2012). According to this model, entrainment would increase gradually during a social episode.

Differentiating between these and other hypotheses is essential in understanding how IBC is achieved and maintained. The main aim of this paper is to present algorithms that will allow researchers to address this. This, we do in section 3. But first, in section 2, we present essential introductory information, starting with an overview of key differences between child and adult EEG that are relevant when conducting hyper scanning research (section 2.1). We then present an overview of different ways of measuring IBC (concurrent vs sequential) and describe how they can be applied to different aspects of the brain signal (amplitude vs power vs phase) (section 2.2).

## Section 2 Identifying different types of connectivity between infant and adult EEG data

### 2.1 Key differences between child and adult EEG

Researchers working with EEG from developmental populations and using naturalistic paradigms also present additional challenges. face several additional challenges as compared to adult EEG researchers using screen-based paradigms (Noreika et al., 2020).

Firstly, due to increased movement within the simulation. This is challenging because signals generated from movement patterns, such as smiling, vocalisations, eye movements, as well as from the neural processing of each of these behaviours, will contribute to the scalp EEG in an unknown way. Although issues of source separation are not new to EEG, it is known that ICA alone fails to separate different sources in data containing high amounts of movement-related activity (e.g. Plöchl et al., 2012; Dimigen, 2020a). This effect is heightened with infant ICA decompositions, which are generally more ambiguous than adult EEG (Marriott Haresign et al., 2021 in press). For example, even simple artifacts such as blink artifacts can be more clearly differentiated from the ongoing EEG in adult data, allegedly because these movements are more stereotypical in adults than in infants.

Secondly, in a traditional, experimenter-designed paradigm, ERPs are calculated relative to experimental events. Although evidence suggests that artifact in traditional experimenter-designed paradigms is still present, and systematically related to experimental events (although see Yuval-Greenberg et al., 2008), the fact that the experiment (and so the artifact) follows a consistent structure means that artifact is relatively easier to deal with. But in a naturalistic paradigm, the events (e.g., eye gaze onsets) are often systematically related to the artifact in the data, and there is no clear and consistent temporal structure. For this kind of data application and optimisation of source separation techniques are crucial (e.g., Dimigen, 2020a). The future study of IBC using naturalistic paradigms will need to control for the contributions of non-neural signal in the EEG. It may also treat this non-neural signal as a data source, by looking at synchrony between these movement-related signals, e.g., synchrony in EMG association with facial affect and vocalisations.

An additional challenge is posed by the intrinsic differences in EEG activity that are observed in recordings from children/ infants compared to adults. For example, we know that the speed at which a brain will process information depends on maturation (e.g., Taylor et al., 2004) and that the canonical frequency bands in child/infant EEG are typically slower than that of adult EEG. For example, peaks in the power density spectrum associated with alpha activity typically observed in the 9-13Hz range in adults can be seen clearly in one-year-old infant EEG between 6 and 9Hz, and are lower still in younger infants (Marshall et al., 2002). This presents a unique problem for developmental researcher’s interested in phase synchrony between infant and adult EEG. One solution to this problem might be to use cross-frequency connectivity methods (Noreika et al., 2020) for example cross-frequency phase coupling (see section 2.2.2.2 for further discussion).

### 2.2 Overview of different ways of measuring inter-brain connectivity (IBC)

IBC can be measured in two ways. First, concurrent IBC (see Figure 1) indexes a zero-lag, simultaneous relationship: ‘at times when A is high, B is also high’ or (for a negative relationship): ‘at times when A is high, B is low’. Concurrent IBC is often referred to using the term ‘synchrony’, and is undirected: A->B is indistinguishable from B->A. Second, sequential IBC indexes a lagged, or temporally oriented relationship: ‘changes in A forward-predict changes in B’. Sequential IBC is directed, and as such, unlike concurrent coupling, it can be asymmetrical: it can be true that A forward predicts B without it being true that B forwards-predicts A, and *vice versa*.

**Figure 1.**
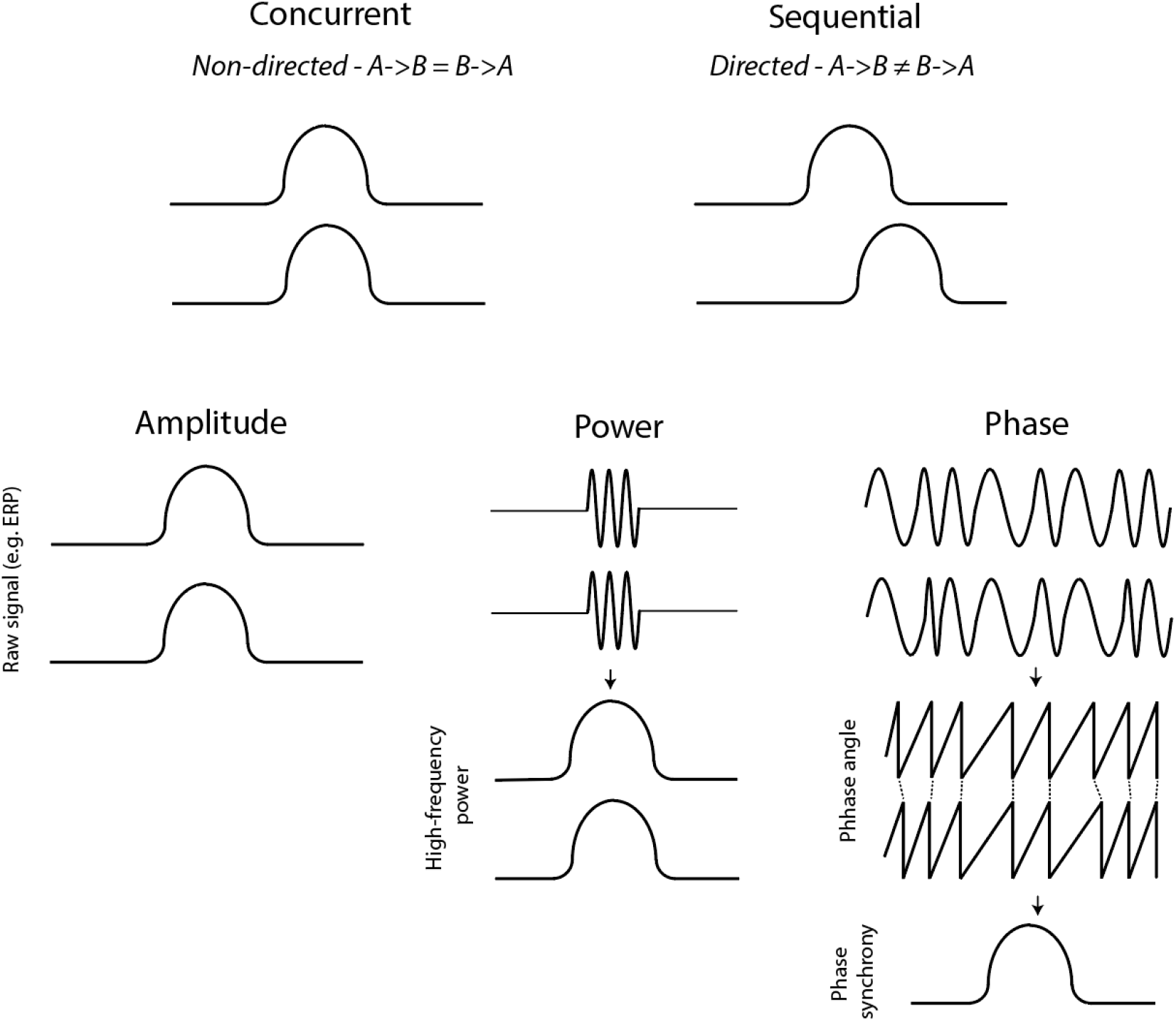
Schematic illustration of the two-connectivity metrics, concurrent and sequential, that we consider in the paper, along with the three aspects of the brain signal: amplitude, power, and phase.

**Figure 2.**
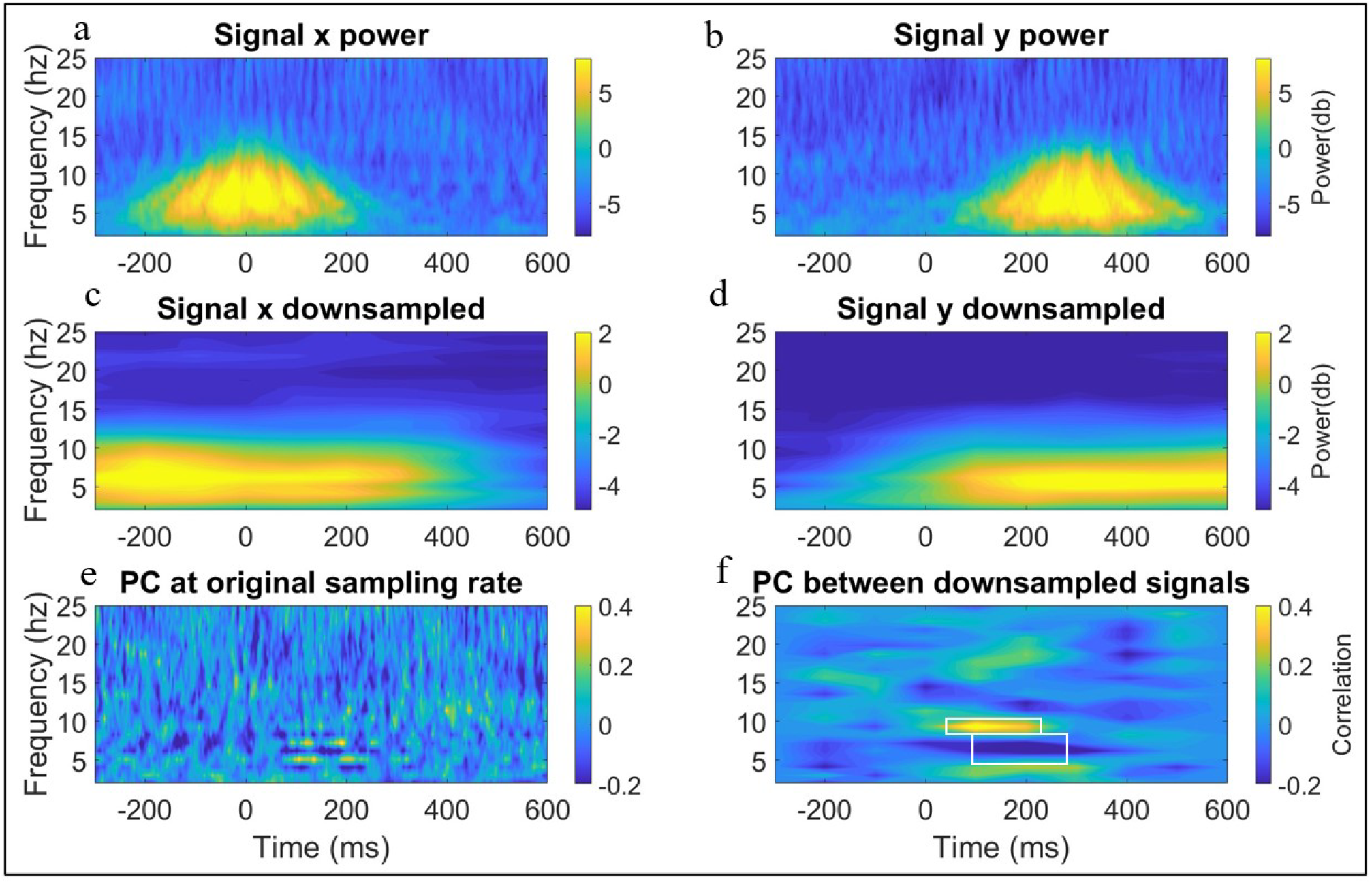
Simulation illustrating the importance of using an appropriate time window for calculating concurrent connectivity. Panel a shows the time-frequency power of signal x. Panel b shows the time-frequency power of signal y. Panel c shows the downsampled (using moving window average) time-frequency power of signal x. Panel d shows the down time-frequency power of signal x. Panel e shows concurrent IBC (spearman’s correlation of single trail power (PC) between x and y) computed at each time-frequency point (i.e., original temporal scale of data). Panel f shows the same concurrent IBC but computed on the downsampled data. The AOI indicates regions of significant correlations.

IBC can also be measured across both the temporal and the frequency domain, and thus across multiple aspects of the brain signal: amplitude, power, and phase (see Figure 1). Currently, most fNIRS and fMRI hyperscanning studies measure co-fluctuations in the amplitude of the signal – which depending on the method, measures blood oxygenation/deoxygenation (fNIRS), the BOLD signal (fMRI) or voltage (for EEG). Measuring IBC as co-fluctuations in the power of the signal at different frequency bands is possible, but not common. Currently, most EEG hyperscanning studies measure phase IBC. There appears, however, to be no reason why these trends should not change in the future.

#### 2.2.1 Measuring concurrent IBC of amplitude and power

Examining concurrent IBC through correlations is one of the simplest and most flexible techniques. Zero-lag concurrent IBC can simply be measured by calculating correlation coefficients between two time series. Spearman’s correlation is generally favoured due to its invariance to non-normally distributed and outlier prone data (Cohen, 2014). The same analysis can apply either to the amplitude of the brain signal or to the power in particular frequency bands. The accompanying code for this section allows the reader to compute single-trial correlations (Spearman’s rho) at each time-frequency point, between pairs of electrodes (e.g., using data from Cz from person one and Cz from person two).

#### 2.2.2 Measuring concurrent IBC of phase – Phase Locking Value (PLV)

Phase locking can be estimated at three levels. First, phase clustering relative to events can be estimated over time and electrodes within a single brain (see SM section 8). Second, phase locking between a single brain and an external stimulus can be calculated over repeated events. Third, phase locking can be estimated at the interpersonal level, between two or more brains. In section 3.1.2 we concentrate on the latter, but two and three are computationally identical. For each, phase locking can be computed at each time-frequency point over trials or in a sliding window within a given trial. This will allow researchers to look at how changes in IBC fluctuate over time. The accompanying code allows for analysis of phase locking within (sliding window) and across trials (for each sample).

PLV, which is the main metric used to estimate phase-locking, measures the extent to which phase angles are similar between two signals over time/trials. PLV is calculated as follows:

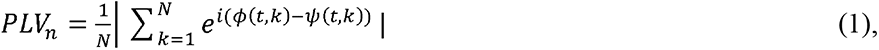

Where N is the number of observations, *ϕ(t, k)* is the phase on observation *k*, at time *t*, in channel *ϕ* and *ψ(t, k)* at channel *ψ*. If the phase angles from the two signals fluctuate over time with a consistent difference, this will lead to PLV values close to 1. If the phase angles fluctuate over time with little consistency between each of the two signals, PLV values will be close to 0. Phase locking measures connectivity between signals with a zero lag. It is worth noting that, as phase synchrony (or phase locking) is a measure of the consistency of phase angles between two signals, where these two cycles are in relation to each other is less important than how they covary.

Any two signals with a common dominant frequency will show relatively consistent variation in phase over time – and hence high phase locking between the signals. This has been used to argue that PLV is particularly prone to detecting spurious hyper connections (Burgess, 2013). However, this is only a problem in a few scenarios. Real EEG data even after narrow-band filtering will still have random variation in the phase of signal over time such that PLV between narrowband filtered signals will also change over time. So, two alpha oscillators will show consistently high PLV only where there is little to no random variation in the phase of the signal over time, which is not a very reasonable assumption for real EEG data.

##### 2.2.2.1 Side note on power and PLV

When analysing any event locked changes in EEG power and/or phase-based connectivity it is important to consider whether these are evoked or induced responses. Evoked responses are additive signals superimposed upon the background/ongoing EEG; induced responses are changes in power and/or phase that happen directly to the background/ongoing EEG. Whilst changes in power/phase resulting from stimulus-locked **evoked** signals could give the appearance of synchronisation, this is interpretationally quite different to potential increases in **induced** neural activity driving increases in connectivity, which is largely what researchers looking at IBC are aiming to measure. For example, if increases in spectral power from two signals are driven by evoked and not induced responses then it is incorrect to examine phase resetting as a potential mechanism behind IBC (or phase-locking more generally) and also incorrect to use the term neural IBC to refer to these mechanisms (e.g., Keitel et al., 2021).

This problem is further complicated because, as Muthukumaraswamy and colleagues (2011) show, transient increases in power can lower error in phase estimation and give the appearance of heightened phase locking (Burgess, 2013). As separating power increases from increases in phase locking is difficult and continually debated (e.g., Sauseng et al., 2007), the best practice for researchers using event-related phase-locking is always to show accompanying power plots.

##### 2.2.2.2 Side note on cross-frequency PLV

As described above (section 2.1) the canonical frequency bands in infant EEG are typically slower compared to adult EEG. It may, therefore, be more appropriate for researchers measuring the quantity of phase-locking between infant and adult EEG to use cross-frequency phase locking. Cross frequency phase synchrony or PLV *m:n* is calculated similarly to PLV as follows:

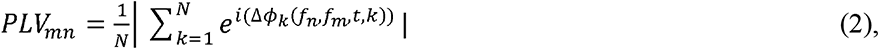

Where, *N* is the number of trials and Δ*ϕ_k_*(*f_n_, f_m_, t, k*) is calculated as follows:

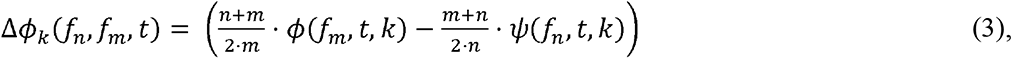

Where *n* and *m* are the centre frequencies of the two signals and should be integer values satisfying the equation *m · f_n_* = *n · f_m_*, and *ϕ*(*f_m_,t*), is the phase angle at channel *ϕ*, at time *t,* on trial *k*, and channel *ψ*. Cross frequency phase synchrony shares the same underlying interpretation as standard phase locking. In accompanying articles in the special edition (Kayhan et al., 2021 in press), we have provided readers with a full pipeline for computing cross-frequency phase-locking on continuous data.

#### 2.2.3 Measuring sequential IBC of amplitude and power – Granger Causality (GC)

The simplest way to measure sequential IBC is simply to repeat the Spearman’s correlation described in section 2.2.1 while shifting one time series forwards or backwards in time relative to the other. For example, if we find that the correlation between two time series x and y is stronger when time series x is backward shifted with respect to time series y, compared with when the simultaneous (‘zero-lag’) correlation between the two-time series is examined, then this indicates that changes in x tend to anticipate changes in y.

Granger Causality is closely related to this approach, but as well as looking at the time-lagged relationship between two time series, it also increases the sensitivity of the prediction by looking at how one time-series forwards predict itself over time (known as the autocorrelation). Given two-time series x and y, Granger Causality is a measure of the extent to which time series x can be predicted by previous samples of y above and beyond how well it can be predicted by previous samples of time series x alone.

GC is defined through the log of ratios of error terms between the bivariate and univariate regressive models, following:

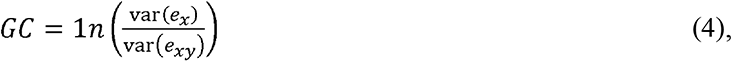

Where *e_x_* is the error term obtained from the univariate autoregressive model fit and *e_xy_* is the error term obtained from the bivariate regressive model fit. Again, the same approach can be adopted to look either at the simple amplitude of the brain signal, or at the power within frequency bands. Time frequency (spectral) GC involves computing the dot product between the regressive coefficients and complex sine waves, analogous to the Fourier transform and then applying those results to the error variance via the transfer function (Cohen, 2014).

Finally, Partially Directed Coherence (PDC) is a frequency domain formulation of GC (Sameshima and Baccala, 2014), measured from the coefficients derived from the autoregressive modelling described above. Although it is not a focus of this paper to compare methods of measuring connectivity, we mention PDC here as it has been used to investigate IBC between adult and infant pairings (e.g., Leong et al 2017; Santamaria, et al 2020). PDC along with other methods of frequency domain connectivity based on autoregressive modelling can be implemented using the extended multivariate autoregressive modelling toolbox (Faes, et al., 2013).

##### 2.2.3.1 Side note on EEG data stationarity and GC

Stationarity refers to whether the statistical properties of data change over time. For example, EEG data that contain low-frequency drifts over time can cause data to become nonstationary. Non-stationarity may take various forms. One form of non-stationarity manifest by unit root processes - where the data may exhibit a stochastic trend or “random drift”. This form of non-stationarity is common in financial time series (e.g., stock prices), but less common in neuroimaging data (because neurophysiological processes are generally physically constrained). Many stationarity tests (including the KPSS test which is implemented within the MVGC toolbox (Barnett and Seth, 2014)) only test for unit-root stationarity. However, unit-root is not the only kind of non-stationarity; there is also “structurally varying” non-stationarity, which may be more common in ERP or more generally task-related data. Here, the statistical properties of the time series change over time, either in a structured/deterministic or stochastically way. (Unit root tests may or may not fail on this form of non-stationarity.) This implies that the Granger causality itself may change over time. Overall stationarity in neuroscience for GC is an ongoing problem (e.g., see Barnett et al., 2018a; 2018b). Current common approaches to address non-stationarity in EEG data include polynomial detrending (Seth et al., 2015), and/or subtraction of the averaged ERP from single-trial data (e.g., Wang et al., 2008) are limited. Another viable solution to address nonstationary EEG data could be to segment the data into shorter time windows in which the data would be stationary enough to perform GC analysis, although this needs more testing.

##### 2.2.3.2 Side note on model order and GC

A crucial parameter to consider when using GC analysis is model order. Model order determines the number of previous samples in the times series that will be used in the autoregressive model fit. For instance, if your data is sampled at 1000hz, a model order of 5 means that the autoregressive model will use a weighted sum of the previous 5ms. The model order used will in part determine the frequency precision of the spectral GC. In our example using only 5ms of data, we would only base our granger prediction on 1/50 of a cycle of 4Hz activity.

To better capture low-frequency dynamics, it can therefore often be useful to down-sample the data prior to analysis. For example, if we resampled our data to 128Hz, still using a model order of 5 would mean we consider the previous 39ms of data in our autoregression. Further considering lower frequency dynamics is a good reason to lean towards the highest model order that is appropriate for the data. In our example increasing the model order to 15 would result in us considering the previous 120ms (at 128Hz), almost half a cycle of 4Hz activity. Some routines exist to help guide the estimation of model order, the most common being Bayes information criterion (BIC) and Akiake information criterion (AIC). Both are implemented within the MVGC toolbox. Importantly, however, what is optimal may vary and different model orders can fit the data equally well.

##### 2.2.3.3 Side note on spectral power and GC

The relationship between spectral power and spectral GC is still uncertain. Research has shown that increases in event-locked spectral power generated from the ERP co-occur with increases in spectral GC (e.g., Wang et al., 2008). It will be important for future research to fully explore the parameters of this relationship (e.g., Winkler et al., 2015) – for example, does the strength of the GC scale linearly with the amount of spectral power? And how is this relationship affected by the sampling rate, signal to noise ratios and so on?

#### 2.2.4 Measuring sequential IBC of phase – Phase Transfer Entropy (PTE)

Phase transfer entropy (PTE) allows researchers to measure sequential IBC of phase. It is calculated using the following equation:

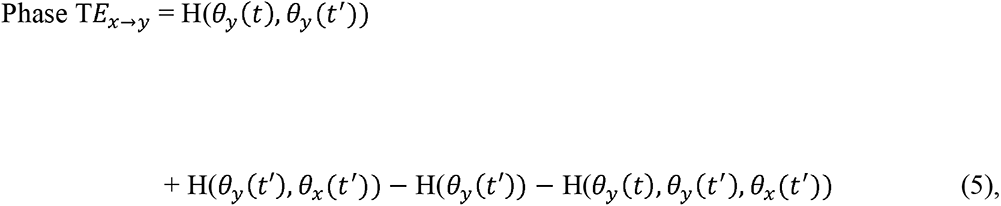

Where *θ*(*t*) is the phase of signal X(t), t’ = t – *δ*, and *θ_x_(t’)* and *θ_y_(t’)* are the previous states of the phase angle time series of x and y, with a given lag of *δ*. Given two-time series x and y, like GC, transfer entropy (TE) estimates whether including the past of x influences our ability to predict y and vice versa. However, unlike GC, TE does this by comparing conditional probabilities (Lobier et al., 2014). E.g., if a signal X ‘causes’/ ‘disambiguates’ a signal Y, then the probability density of the future of Y conditioned on its past should be different from the probability density of the future of Y conditioned on the pasts of both X and Y. As transfer entropy is based on the same underlying principles as GC, it has been shown that results obtained using GC and PTE are identical for Gaussian variables (e.g. Barnett et al., 2009). Therefore, results from phase transfer entropy analyses are therefore similar to results obtained from GC analysis but based on the phase of the time series. If there is an increase in PTE between two time series, then we can say that there has been some information transfer from one phase angle time series to the other.

Whilst Phase Transfer Entropy has not been widely used within cognitive neuroscience as a framework for analysing connectivity patterns between two systems, it has many advantages and useful properties. For example, as entropy is not based on the temporal structure of the data, it can be computed over time and trials - whereas other measures e.g., PLV can only be computed over time or trials, not both. This is a major advantage as including data from time and trials simultaneously means that entropy can be computed in shorter time windows than other window-based connectivity measures (e.g., GC), thus retaining a greater degree of the original temporal precision of the data whilst still having sufficiently high signal to noise ratios (Cohen, 2014).

### 2.3 Cautionary note on importance of temporal scale

In this section we highlight how temporal scale influences concurrent IBC. To illustrate this, we simulated two signals that show an event-locked transient increase in spectral power which peaks 300ms later in signal y compared with signal x (Figure 3a, 3b). Details of the time-frequency decomposition can be found in SM 7. To compute concurrent IBC, we performed two calculations: first, we calculated Spearman’s correlations between the power of time series x and y at each time-frequency point independently (see Fig 3e). Second, we first downsampled the data using a 0.5s sliding window with 200ms of overlap between successive windows (Figs 3c, 3d) before repeating the same analysis (Fig 3f). When using a fine temporal scale, we detect no changes in concurrent IBC, but when using a larger time window (reduced temporal precision) we do (Fig 3f). This illustrates how the pre-processing of data prior to IBC analyses can alter the results.

**Figure 3.**
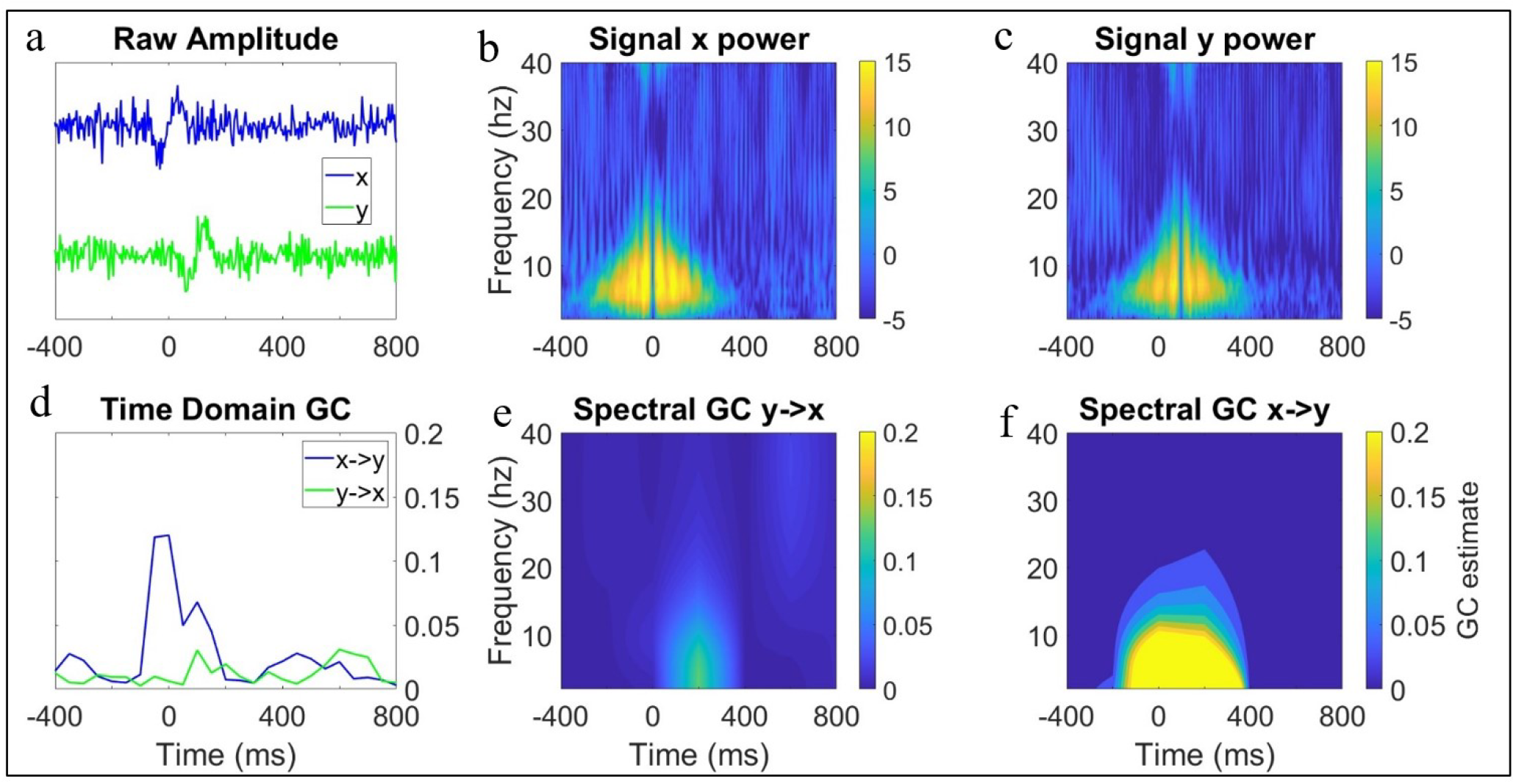
Simulated data showing different mechanisms that that could give rise to increases in interbrain granger causality between parents and infants. (a) shows the two correlated (single-trial amplitude) transient signals x and y. Y was generated from previous samples of x with a lag of 100ms, such that (d) there is a substantial event locked increase in GC from x to y but no GC influence from y to x. (b) shows time-frequency power from signal x from panel a. (c) shows time-frequency power from signal y from panel a. (e) shows spectral GC from y to x and (g) from x to y.

## Section 3 Differentiating event-locked from non-event-locked changes in IBC

In this section we present algorithms and methods to allow researchers to compute the metrics described above. We also present simulations in which we artificially introduce a given relationship between two time series, in order to assess how well the algorithms detect it. Throughout the section, we also discuss how different signal changes can manifest either as transient, event-locked changes (section 3.1) or as slower, more continuous, non-event-locked changes (section 3.2). In section 3.3, we describe methods to quantify whether observed event-locked changes in IBC differed significantly from chance.

### 3.1 Simulations - event-locked changes

#### 3.1.1 Amplitude and power

At its most simple level, computing the correlation coefficients for single trail time-frequency amplitude/ power does not involve of any algorithms more complex than a Spearman’s correlation. Although we have not focussed on it here, we do still provide routines for computing concurrent amplitude/ power connectivity in the accompanying code.

To illustrate how event-locked neural responses might give rise to changes in sequential amplitude/power IBC we simulated two ERP-like signals (x and y) (see Figure 3). Signal y was generated from previous samples of x plus noise (see SM sections 2 and 7 for full details). From this simulation, it can be seen that the sequential IBC between x and y observable in the raw data (Fig 3a) manifests as strong x->y GC influences but not y->x GC influences, as expected (Fig 3a). When the same analysis is applied to the power of the signal (Fig 3b, 3c), the predicted results are again observed. Spectral x->y GC influences are observed across a range of lower frequencies (Fig 3e), but no spectral y->x effects are observed (Fig 3f).

In the accompanying code for this section, we also provide the user with code for measuring Spearman’s correlations of spectral power between two neural responses, and implementation of time domain and spectral GC (i.e., sequential IBC based on amplitude/power) for measuring event locked changes in dual EEG data. The user will also be able to easily specify more advanced parameters such as the time window size and model order used for the time-varying GC estimates.

#### 3.1.2 Phase

To illustrate how event-locked neural responses might give rise to changes in concurrent phase-based IBC, we simulated two partially phase-locked signals (x and y) with a concurrent phase reset/modulation +200ms after an event (time 0) (see Fig 4a) (see SM section 4 for more details). From this simulation, it can be seen that, during the time window following the manipulation at +200ms, the phase angles of the two time series converge (Fig 4a) as expected. The phase synchronisation values of the two time series also converge (Fig 4c) as expected.

**Figure 4.**
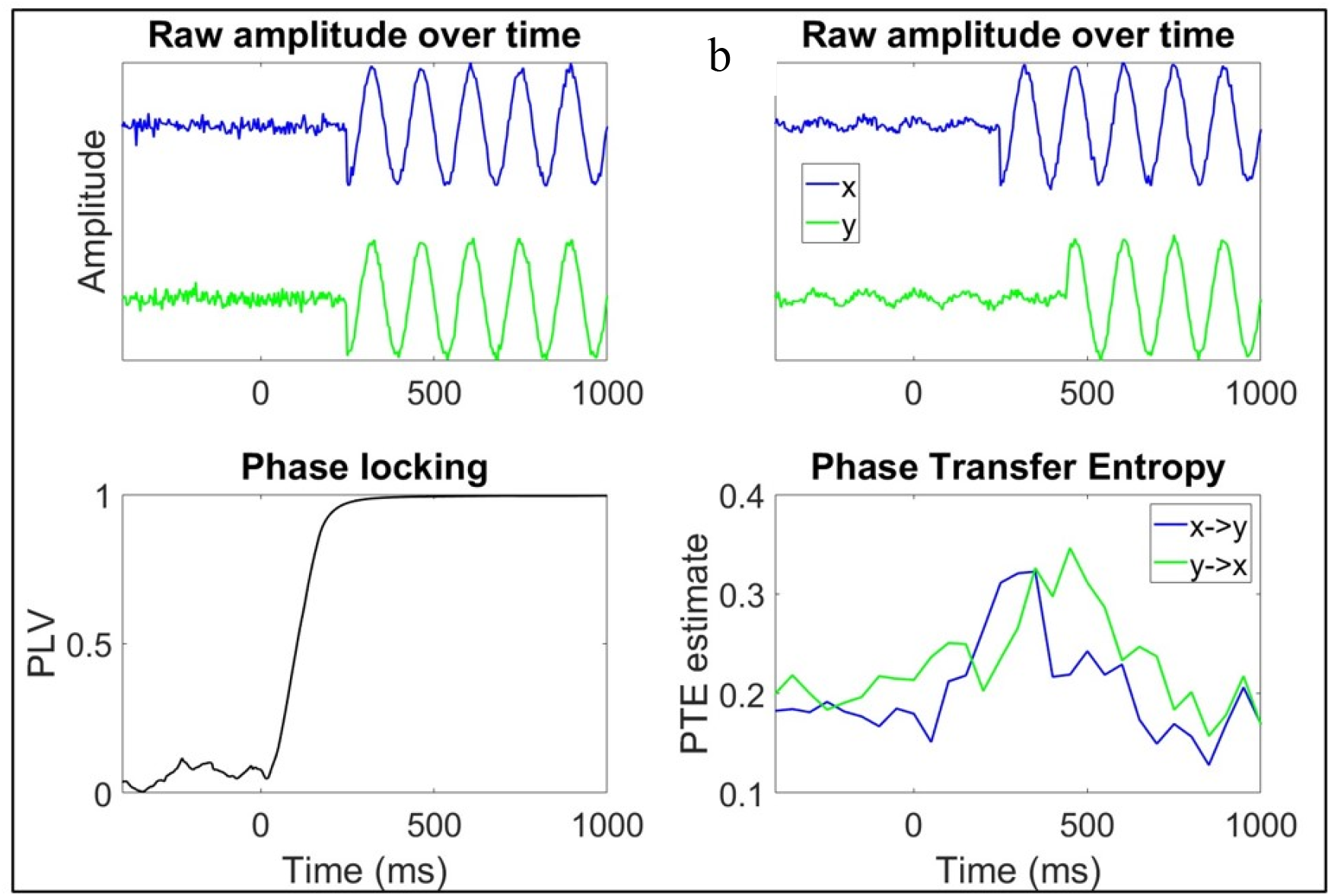
Simulated data showing how event locked phase modulations could give rise to inter brain phase-based IBC between parents and infants. (a) Time series data x and y were subjected to a phase reset at +200ms and become purely phase-locked. The increased consistency in phase angle circa +200ms between x and y yields (c) a notable increase in the phase-locking between x and y ∼200ms. (b) simulated data showing a situation in which the phase modulations in one signal (x) predict the phase modulations in another signal (y) and how this lagged/ directed relationship in phase can be captured (d) using phase transfer entropy.

To illustrate the sequential phase IBC, we simulated x and y again in the same way but here the phase modulation in y occurred 200ms later than in signal x (see Fig 4b) (see SM section 5 for more details). From this simulation it can be seen that when phase modulations in one signal occur later/ earlier than phase modulations in another signal (e.g., x in Fig 4b becomes phase-locked at +200ms, and y in Fig 4b becomes phase-locked at 400ms) and that these modulations are correlated, then this relationship (or form of sequential IBC) can be captured using directed phase IBC methods such as phase transfer entropy – illustrated by the increase in PTE in the time window between phase modulations of x and y (∼+200-400ms) (Fig 4d).

In the accompanying code for this section, we provide the user with full implementations of inter-individual, time-frequency PLV and PTE (i.e., sequential IBC based on phase). The user can easily specify parameters such as time window size for PLV/PTE as well as the model order for PTE.

### 3.2 Simulation – non-event-locked changes

#### 3.2.1 Amplitude and power

To illustrate how gradual changes in amplitude/power IBC that are not time-locked to the onset of an event might arise, we simulated two oscillatory signals (x and y) where y was generated from previous samples of x plus noise (see Fig 5a). To simulate a gradual change in GC we reduced this noise parameter over time (see SM section 3 and 7 for more details). From this simulation, it can be seen that, as expected, the x->y GC influences increase during the time window, but no changes in y->x GC influences are observed (Fig 5b).

**Figure 5.**
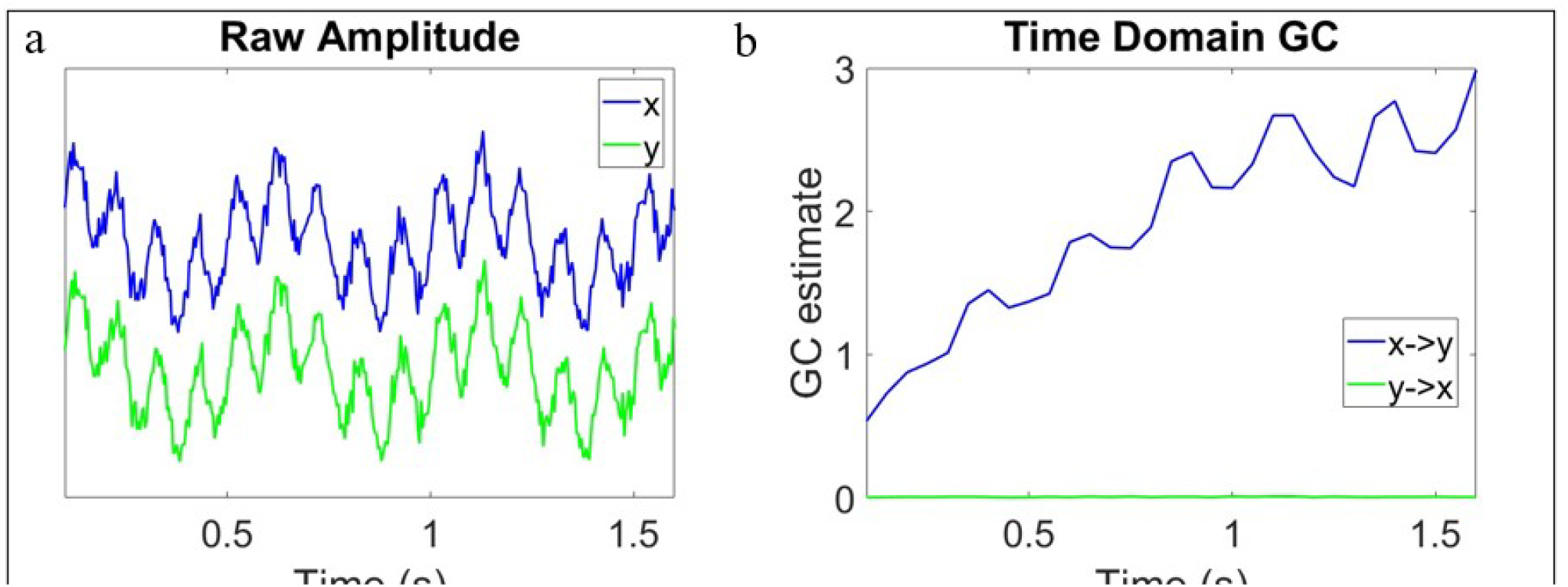
Simulated data showing different mechanisms that that could give rise to increases in interbrain granger causality between parents and infants. (a) shows two oscillatory signals. Y was generated as a product of previous samples of x with a lag of 25ms. We decreased the amount of noise in x over time to simulate (b) a gradual increase in GC from x to y throughout the segment

In the accompanying code for this section we show how the same code from the previous section (3.1.1) can be leveraged to look at questions regarding non-event locked changes in inter brain IBC. No new algorithms are implemented here.

#### 3.2.2 Phase

To illustrate how gradual changes in phase IBC might arise that are not time-locked to the onset of an event, we simulated two oscillatory signals (x and y) with slow drifts in peak frequency over time (signal x linearly increased in peak frequency from 6 to 9hz and signal y decreased from 12-9hz) (see Figure 6). Full details of how we simulated this data and the time-frequency decomposition can be found in the supplementary materials (SM 6 and 7). From this simulation, it can be seen that the closer the signals become in peak frequency the more consistent the relationship between phase angles over time is, and thus the higher the phase-locking values between x and y is.

**Figure 6.**
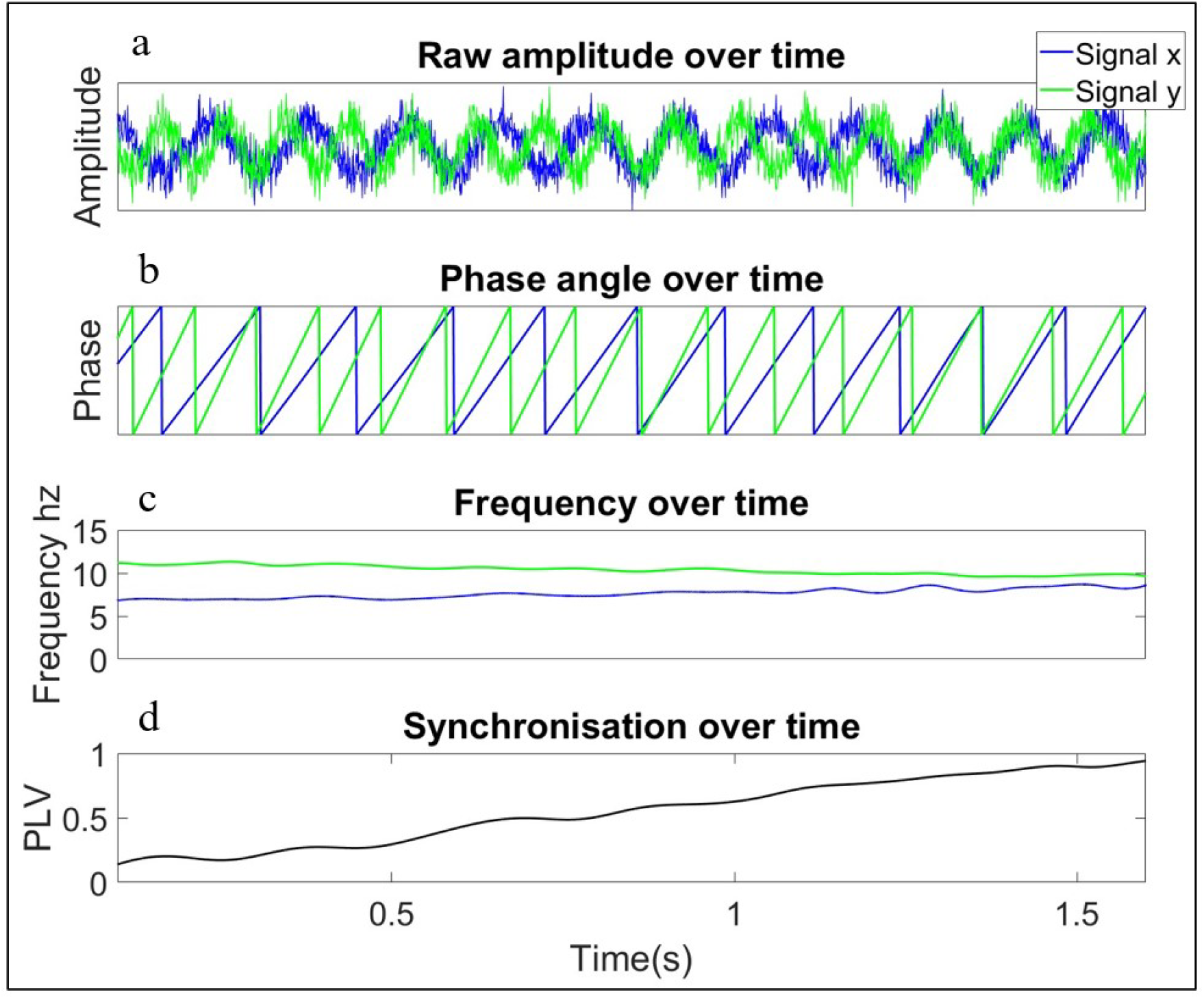
Simulated data showing the different mechanisms that could give rise to increases interbrain phase locking between parents and infants. (a) Time series data x and y both exhibit slow trends in frequency over time toward a common dominant frequency. (c) signal x increases from 6 to 9hz over the time course whereas time series y decreases from 12 to 9hz. The closer the signal become in peak frequency (b) the more consistent the relationship between phase angles over time is, (d) yielding a gradual increase in PLV between x and y over time

In the accompanying code for this section, we show how the same code from the previous section (3.1.2) can be leveraged to look at questions regarding non-event locked changes in inter brain IBC. No new algorithms are implemented here.

### 3.3 Quantifying event-locked changes

In this final section we consider the question of statistical significance. If we know that specific events occurred in the data, how can we test whether statistically significant changes in IBC occurred relative to these events?

#### 3.3.1 Amplitude and power

Significant changes in GC can be evaluated with an F-statistic (Barnett & Seth, 2014), which can be implemented through the MVGC toolbox. Statistical significance can also be obtained via permutation testing, which can be applied to power correlations, time domain and spectral GC (Maris & Oostenveld, 2007). For measuring changes in GC that are strongly time/event locked, permuting the order of the time segments within trials is generally recommended over permuting the trial order whilst leaving the time segments intact (Cohen, 2014).

#### 3.3.2 Phase

Assuming a von Mises distribution (normal distribution for circular data) statistical significance of ITC and PLV can be evaluated against a p-value, approximated against the null hypothesis using the Rayleighs test which can be implemented using the Circstat toolbox (Berens, 2009). Statistical significance can also be assessed against a threshold ITC/PLV value (Cohen, 2014). Any values which exceed this resulting threshold can be considered significant. Alternatively, the significance of time-frequency varying ITC and PLV as well as PTE (when computed in a sliding window within trials) can also be assessed using nonparametric permutation testing (e.g., Maris & Oostenveld, 2007)

## Section 4 limitations of event locked analysis for studying inter brain neural synchrony

*Procambarus clarkia*, a breed of freshwater crayfish, exhibit only a small range of social behaviours, primarily focussed around dominance/ subordinate, yet their physiological systems are capable of supporting these interactions as well as intra interactions with their environment with a remarkable level of temporal fidelity (e.g Schapker et al., 2002). The way that humans interact with their environment and each other is infinitely more complex and multi-layered (Hasson & Frith, 2016; Hoehl et al., 2021; Murray et al., 2016). However, most researchers who study interacting physiological systems during social engagement typically do so using methods that show how connectivity varies by topography and between different experimental conditions (or participants), but which obscures how connectivity varies over time. In this article, we have argued that this omission blocks us from developing a mechanistic understanding of how connectivity is established and maintained.

In this article, we presented algorithms that allow researchers to measure how connectivity between two physiological systems varies as a function of time. We have differentiated between two types of connectivity: concurrent (‘when A is high, B is high’) and sequential entrainment (‘changes in A forward-predict changes in B’) (section 2.2). And we have described how these measures can be applied to three aspects of the neural signal: amplitude, power, and phase (section 2.2, see Figure 1).

Concurrent connectivity of amplitude and power can be measured using correlations (section 2.2.1). Measuring concurrent connectivity in phase is less straightforward (section 2.2.2): it is measured using Phase-Locking Value (PLV), but challenges remain in understanding the relationship between power and PLV (2.2.2.1), and in understanding how intrinsic differences between children and adults in the canonical frequency bands influence child-adult PLV (2.2.2.2).

Sequential connectivity of amplitude and power can be measured using Granger Causality (GC), which can be applied to look either at the amplitude of the brain signal or the power within frequency bands (section 2.2.3). Again, several challenges exist in applying analyses using GC to EEG data, such as the problem of ensuring that data is stationary (section 2.2.3.1), the importance of model order in influencing the results of GC calculations (section 2.2.3.2) and the relationship between spectral power and GC (section 2.2.3.3). Sequential connectivity of phase can be measured using Phase Transfer Entropy (section 2.2.4). Although not widely used in neuroimaging research, we have argued that this approach has promise.

In section 3 we also presented algorithms for measuring connectivity changes relative to event-locked (section 3.1) and non-event-locked (section 3.2) changes, along with techniques for exploring whether observed event-locked changes differ significantly from chance (section 3.3). These algorithms are illustrated using simulated data in which different types of relationship are artificially introduced into the data, to allow the reader to see how the algorithm detects them. These simulations also generated some useful insights into how, for example, gradual alignment in the dominant power of oscillatory activity across two systems can lead to increases in Phase Locking Value (section 3.2.2).

### 4. Limitations

There are, of course, many limitations to this work. The field of inter-brain connectivity using EEG is still in its infancy, and substantial questions remain about how satisfactorily artefact can be removed from brain data (section 2.1); in understanding the relationship between power changes and IBC (sections 2.2.2.1 and 2.2.3.3) and the influence of model order on Granger Causal analyses (section 2.2.3.2); and so on.

Two further limitations should be noted. First, we are often considering events that have different periodic structures, and that unfold over different time scales. For example, researchers might want to examine the relationship between eye gaze shifts (which take place every ∼300ms – i.e. at ∼3Hz), changes in autonomic arousal (between ∼0.01and ∼0.5Hz) and changes in EEG (between ∼2Hz-∼30Hz). Although we have presented some methods for looking at this – such as temporal correlations in power at different frequencies (section 2.2.1) and cross-frequency PLV (section 2.2.2.2) – we have not discussed other approaches, such as phase-amplitude coupling (Canolty & Knight, 2010; Tort et al., 2010) that would also be useful.

Second, we have concentrated exclusivity on bivariate connectivity. Unlike the ways in which social information processing is typically studied (i.e., using repeated, discrete and unecological screen-based stimulus), real social interactions involve highly layered and complex sequences of multimodal events that can unfold over multiple time scales in a continuous and interdependent way. For example, consider the multimodal pathways to joint attention as illustrated by Yu and Smith (2016), in which often sequences of social interactions between parents and infants can involve initiating and responding to various postural and gestural movements, as well as visual (gaze) information and vocalisations, presented in combination. Future work will require more advanced data analytics, and the collection of larger datasets, to address this.

#### 4.3 Conclusion

Despite these limitations, the present focus on fine-grained neural responses has the potential to provide valuable new insights into the cognitive/ attention processes that support dynamic social interaction, beyond what is possible using current/ standard approaches in hyper scanning. Arguably, these limitations and considerations reinforce the idea that analysing the rich temporal dynamics of neural activity brings us closer to the true complexity of brain function.

## Implementation

All MATLAB functions for the implementation of the various algorithms presented here for event locked connectivity analysis of dual EEG data, including a sample dual EEG dataset, as well as, code for generating the simulated data can be found here [https://github.com/Ira-marriott/Using-dual-EEG-to-analyse-event-locked-changes-in-child-adult-neural-entrainment-.git]

## Acknowledgements

This research was funded by Project Grant RPG-2018-281 from the Leverhulme Trust, by an ERC Marie Curie Fellowship JDIL 845859 and by ERC grant number ONACSA 853251. Thanks to members of the UEL BabyDev Lab, Mike X Cohen and Lionel Barnett for useful advice.

## References

Barnett, L., Barrett, A. B., & Seth, A. K. (2009). Granger causality and transfer entropy are equivalent for Gaussian variables. Physical review letters, 103(23), 238701.

Barnett, L., Barrett, A. B., & Seth, A. K. (2018). Misunderstandings regarding the application of Granger causality in neuroscience. Proceedings of the National Academy of Sciences, 201714497

Barnett, L., Barrett, A. B., & Seth, A. K. (2018). Solved problems for Granger causality in neuroscience: A response to Stokes and Purdon. NeuroImage, 178, 744–748.

Barnett, L., & Seth, A. K. (2014). The MVGC multivariate Granger causality toolbox: a new approach to Granger-causal inference. Journal of neuroscience methods, 223, 50–68.

Berens, P. (2009). CircStat: a MATLAB toolbox for circular statistics. J Stat Softw, 31(10), 1–21.

Bloom, K. (1988). Quality of adult vocalizations affects the quality of infant vocalizations. Journal of child language, 15(3), 469–480.

Bögels, S. (2020). Neural correlates of turn-taking in the wild: Response planning starts early in free interviews. Cognition, 203, 104347.

Burgess, A. P. (2012). Towards a unified understanding of event-related changes in the EEG: the firefly model of synchronization through cross-frequency phase modulation. PloS one, 7(9), e45630.

Burgess, A. P. (2013). On the interpretation of synchronization in EEG hyperscanning studies: a cautionary note. Frontiers in human neuroscience, 7, 881.

Canolty, R. T., & Knight, R. T. (2010). The functional role of cross-frequency coupling. Trends in cognitive sciences, 14(11), 506–515

Cohen, M. X. (2014). Analyzing neural time series data: theory and practice. MIT press.

Dimigen, O. (2020). Optimizing the ICA-based removal of ocular EEG artifacts from free viewing experiments. NeuroImage, 207, 116117.

Dumas, G., Nadel, J., Soussignan, R., Martinerie, J., & Garnero, L. (2010). Inter-Brain Synchronization during Social Interaction. PLoS ONE, 5(8), e12166.

Faes, L., Erla, S., Porta, A., & Nollo, G. (2013). A framework for assessing frequency domain causality in physiological time series with instantaneous effects. Philosophical Transactions of the Royal Society A: Mathematical, Physical and Engineering Sciences, 371(1997), 20110618.

Farroni, T., Csibra, G., Simion, F., & Johnson, M. H. (2002). Eye contact detection in humans from birth. Proceedings of the National academy of sciences, 99(14), 9602–9605.

Golumbic, E. M. Z., Ding, N., Bickel, S., Lakatos, P., Schevon, C. A., McKhann, G. M., … & Schroeder, C. E. (2013). Mechanisms underlying selective neuronal tracking of attended speech at a “cocktail party”. Neuron, 77(5), 980–991.

Giraud, A. L., & Poeppel, D. (2012). Cortical oscillations and speech processing: emerging computational principles and operations. Nature neuroscience, 15(4), 511

Grossmann, T., Johnson, M. H., Farroni, T., & Csibra, G. (2007). Social perception in the infant brain: gamma oscillatory activity in response to eye gaze. Social cognitive and affective neuroscience, 2(4), 284–291.

Haresign, I. M., Phillips, E., Whitehorn, M., Noreika, V., Jones, E., Leong, V., & Wass, S. V. (2021). Automatic classification of ICA components from infant EEG using MARA. bioRxiv.

Hamilton, A. F. D. C. (2021). Hyperscanning: beyond the hype. Neuron, 109(3), 404–407.

Hasson, U., & Frith, C. D. (2016). Mirroring and beyond: coupled dynamics as a generalized framework for modelling social interactions. Philosophical Transactions of the Royal Society B: Biological Sciences, 371(1693), 20150366.

Hirsch, J., Zhang, X., Noah, J. A., & Ono, Y. (2017). Frontal temporal and parietal systems synchronize within and across brains during live eye-to-eye contact. NeuroImage, 157, 314–330.

Hoehl, S., Fairhurst, M., & Schirmer, A. (2021). Interactional synchrony: signals, mechanisms and benefits. Social Cognitive and Affective Neuroscience, 16(1-2), 5–18.

Hoehl, S., & Striano, T. (2008). Neural processing of eye gaze and threat□related emotional facial expressions in infancy. Child development, 79(6), 1752–1760.

Hoehl, S., & Striano, T. (2010). Infants’ neural processing of positive emotion and eye gaze. Social Neuroscience, 5(1), 30–39.

Jones, E. J. H., Goodwin, A., Orekhova, E., Charman, T., Dawson, G., Webb, S. J., & Johnson, M. H. (2020). Infant EEG theta modulation predicts childhood intelligence. Scientific reports, 10(1), 1–10.

Keitel, C., Obleser, J., Jessen, S., & Henry, M. J. (2021). Frequency-Specific Effects in Infant Electroencephalograms Do Not Require Entrained Neural Oscillations: A Commentary on Köster et al.(2019). Psychological Science, 09567976211001317.

Kingsbury, L., Huang, S., Wang, J., Gu, K., Golshani, P., Wu, Y. E., & Hong, W. (2019). Correlated Neural Activity and Encoding of Behavior across Brains of Socially Interacting Animals. Cell.

Kinreich, S., Djalovski, A., Kraus, L., Louzoun, Y., & Feldman, R. (2017). Brain-to-brain synchrony during naturalistic social interactions. Scientific reports, 7(1), 1–12.

Kirkland, A. (2020). An exploration of the neural correlates of turn-taking in spontaneous conversation.

Kuhl, P. K., Tsao, F. M., & Liu, H. M. (2003). Foreign-language experience in infancy: Effects of short-term exposure and social interaction on phonetic learning. Proceedings of the National Academy of Sciences, 100(15), 9096–9101.

Lachat, F., & George, N. (2012). Oscillatory brain correlates of live joint attention: a dual-EEG study. Frontiers in human neuroscience, 6, 156.

Leong, V., Byrne, E., Clackson, K., Georgieva, S., Lam, S., & Wass, S. (2017). Speaker gaze increases information coupling between infant and adult brains. Proceedings of the National Academy of Sciences, 114(50), 13290–13295.

Leong, V., Noreika, V., Clackson, K., Georgieva, S., Brightman, L., Nutbrown, R., . . . Wass, S. (2019). Mother-infant interpersonal neural connectivity predicts infants’ social learning.

Liu, D., Liu, S., Liu, X., Zhang, C., Li, A., Jin, C., . . . Zhang, X. (2018). Interactive Brain Activity: Review and Progress on EEG-Based Hyperscanning in Social Interactions. Frontiers in Psychology, 9, 1862.

Lobier, M., Siebenhühner, F., Palva, S., & Palva, J. M. (2014). Phase transfer entropy: a novel phase-based measure for directed connectivity in networks coupled by oscillatory interactions. Neuroimage, 85, 853–872.

Makeig, S., Debener, S., Onton, J., & Delorme, A. (2004). Mining event-related brain dynamics. Trends in cognitive sciences, 8(5), 204–210

Mandel, A., Bourguignon, M., Parkkonen, L., & Hari, R. (2016). Sensorimotor activation related to speaker vs. listener role during natural conversation. Neuroscience letters, 614, 99–104.

Maris, E., & Oostenveld, R. (2007). Nonparametric statistical testing of EEG-and MEG-data. Journal of neuroscience methods, 164(1), 177–190.

Moreau, Q., & Dumas, G. (2021). Beyond “correlation vs. causation”: multi-brain neuroscience needs explanation. Trends Cogn. Sci. Published online March, 19, 2021.

Müller, V., & Lindenberger, U. (2019). Dynamic Orchestration of Brains and Instruments During Free Guitar Improvisation. Frontiers in integrative neuroscience, 13, 50.

Murray, L., De Pascalis, L., Bozicevic, L., Hawkins, L., Sclafani, V., & Ferrari, P. F. (2016). The functional architecture of mother-infant communication, and the development of infant social expressiveness in the first two months. Scientific Reports, 6(1), 1–9

Muthukumaraswamy, Suresh D., and Krish D. Singh. “A cautionary note on the interpretation of phase-locking estimates with concurrent changes in power.” Clinical neurophysiology 122.11 (2011): 2324–2325.

Noreika, V., Georgieva, S., Wass, S., & Leong, V. (2020). 14 challenges and their solutions for conducting social neuroscience and longitudinal EEG research with infants. Infant Behavior and Development, 58, 101393.

Novembre, G., & Iannetti, G. D. (2020). Hyperscanning Alone Cannot Prove Causality. Multibrain Stimulation Can. Trends in Cognitive Sciences.

Nguyen, T., Schleihauf, H., Kayhan, E., Matthes, D., Vrtička, P., & Hoehl, S. (2020). The effects of interaction quality on neural synchrony during mother-child problem solving. cortex, 124, 235–249.

Nguyen, T., Schleihauf, H., Kayhan, E., Matthes, D., Vrtička, P., & Hoehl, S. (2021). Neural synchrony in mother–child conversation: Exploring the role of conversation patterns. Social Cognitive and Affective Neuroscience, 16(1-2), 93–102.

Plöchl, M., Ossandón, J. P., & König, P. (2012). Combining EEG and eye tracking: identification, characterization, and correction of eye movement artifacts in electroencephalographic data. Frontiers in human neuroscience, 6, 278.

Piazza, E. A., Cohen, A., Trach, J., & Lew-Williams, C. (2021). Neural synchrony predicts children’s learning of novel words. Cognition, 214, 104752.

Pérez, A., Carreiras, M., & Duñabeitia, J. A. (2017). Brain-to-brain entrainment: EEG interbrain synchronization while speaking and listening. Scientific reports, 7(1), 1–12.

Pérez, A., Dumas, G., Karadag, M., & Duñabeitia, J. A. (2019). Differential brain-to-brain entrainment while speaking and listening in native and foreign languages. Cortex, 111, 303–315.

Quadrelli, E., Conte, S., Macchi Cassia, V., & Turati, C. (2019). Emotion in motion: Facial dynamics affect infants’ neural processing of emotions. Developmental psychobiology, 61(6), 843–858

Redcay, E., & Warnell, K. R. (2018). A social-interactive neuroscience approach to understanding the developing brain. Advances in child development and behavior, 54, 1–44.

Redcay, E., & Schilbach, L. (2019). Using second-person neuroscience to elucidate the mechanisms of social interaction. Nature Reviews Neuroscience, 20(8), 495–505

Reindl, V., Gerloff, C., Scharke, W., & Konrad, K. (2018). Brain-to-brain synchrony in parent-child dyads and the relationship with emotion regulation revealed by fNIRS-based hyperscanning. NeuroImage, 178, 493–502.

Reissland, N., & Stephenson, T. (1999). Turn□taking in early vocal interaction: a comparison of premature and term infants’ vocal interaction with their mothers. Child: care, health and development, 25(6), 447–456.

Reindl, V., Wass, S., Leong, V., Scharke, W., Wistuba, S., Wirth, C. L., … & Gerloff, C. (2021). Synchrony of mind and body are distinct in mother-child dyads. bioRxiv.

Samaha, J., & Postle, B. R. (2015). The speed of alpha-band oscillations predicts the temporal resolution of visual perception. Current Biology, 25(22), 2985–2990.

Sameshima, K., & Baccala, L. A. (Eds.). (2014). Methods in brain connectivity inference through multivariate time series analysis. CRC press.

Santamaria, L., Noreika, V., Georgieva, S., Clackson, K., Wass, S., & Leong, V. (2020). Emotional valence modulates the topology of the parent-infant inter-brain network. NeuroImage, 207, 116341.

Sauseng, P., Klimesch, W., Gruber, W. R., Hanslmayr, S., Freunberger, R., & Doppelmayr, M. (2007). Are event-related potential components generated by phase resetting of brain oscillations? A critical discussion. Neuroscience, 146(4), 1435–1444.

Seth, A. K., Barrett, A. B., & Barnett, L. (2015). Granger causality analysis in neuroscience and neuroimaging. Journal of Neuroscience, 35(8), 3293–3297.

Schapker, H., Breithaupt, T., Shuranova, Z., Burmistrov, Y., & Cooper, R. L. (2002). Heart and ventilatory measures in crayfish during environmental disturbances and social interactions. Comparative Biochemistry and Physiology Part A: Molecular & Integrative Physiology, 131(2), 397–407.

Schilbach, L., Timmermans, B., Reddy, V., Costall, A., Bente, G., Schlicht, T., & Vogeley, K. (2013). Toward a second-person neuroscience. Behavioral and brain sciences, 36(4), 393–414.

Simony, E., Honey, C. J., Chen, J., Lositsky, O., Yeshurun, Y., Wiesel, A., & Hasson, U. (2016). Dynamic reconfiguration of the default mode network during narrative comprehension. Nature communications, 7, 12141.

Taylor, M. J., Batty, M., & Itier, R. J. (2004). The faces of development: a review of early face processing over childhood. Journal of cognitive neuroscience, 16(8), 1426–1442.

Tort, A. B., Komorowski, R., Eichenbaum, H., & Kopell, N. (2010). Measuring phase-amplitude coupling between neuronal oscillations of different frequencies. Journal of neurophysiology, 104(2), 1195–1210.

Wang, X., Chen, Y., & Ding, M. (2008). Estimating Granger causality after stimulus onset: a cautionary note. Neuroimage, 41(3), 767–776.

Wass, S. V., Noreika, V., Georgieva, S., Clackson, K., Brightman, L., Nutbrown, R., Santamaria, L., & Leong, V. (2018). Parental neural responsivity to infants’ visual attention: how mature brains influence immature brains during social interaction. PLoS biology, 16(12), e2006328

Wass, S. V., Whitehorn, M., Haresign, I. M., Phillips, E., & Leong, V. (2020). Interpersonal neural entrainment during early social interaction. Trends in cognitive sciences, 24(4), 329–342.

Wilson, G. T. (1972). The factorization of matricial spectral densities. SIAM Journal on Applied Mathematics, 23(4), 420–426.

Winkler, I., Haufe, S., Porbadnigk, A. K., Müller, K. R., & Dähne, S. (2015). Identifying Granger causal relationships between neural power dynamics and variables of interest. NeuroImage, 111, 489–504.

Wutz, A., Melcher, D., & Samaha, J. (2018). Frequency modulation of neural oscillations according to visual task demands. Proceedings of the National Academy of Sciences, 115(6), 1346–1351.

Yu, C. and L.B. Smith, The social origins of sustained attention in one-year-old human infants. Current Biology, 2016. 26(9): p. 1235–1240.

Yuval-Greenberg, S., Tomer, O., Keren, A. S., Nelken, I., & Deouell, L. Y. (2008). Transient induced gamma-band response in EEG as a manifestation of miniature saccades. Neuron, 58(3), 429–441.

Zhang, W., & Yartsev, M. M. (2019). Correlated Neural Activity across the Brains of Socially Interacting Bats. Cell.

