## Supplementary materials for "Using dual EEG to analyse event-locked changes in child-adult neural connectivity"

**Supporting Materials**

**Appendix A**

1. *Simulation for event locked changes in concurrent amplitude/ power connectivity*

To illustrate how event locked changes might give rise to changes in concurrent amplitude/power inter brain connectivity (IBC) (see section 2.3 of main text), we simulated two time series x and y each embedded with one purely phase-locked transient oscillation as a product of a sine wave (640 samples, at a sampling rate of 256 Hz) at 7Hz convolved with a gaussian kernel (640 samples, sd of 0.1). The peak of the transient oscillation of time series y was offset from the peak of time series x by +100ms. This was done to be practically most similar to the examples on measuring changes in sequential amplitude/ power based connectivity (e.g., see figure 3 of main text), but to still highlight how changes in event locked concurrent power connectivity can be measured at a fine temporal scale (e.g., looking at sub-second changes). We simulated 100 trials of data in this way. Details on how we derived single trial EEG power can be found in section 7. Spearman's correlation was calculated between signals x and y at each time-frequency point.

2. *Simulation for event locked changes in sequential amplitude/ power connectivity*

To illustrate how event-locked neural responses might give rise to changes in sequential amplitude/power IBC (see section 3.1.1. of main text), we simulated two time series x and y each embedded with one transient oscillation as a product of a sine wave (640 time points, at a sampling rate of 256 Hz) at 7Hz convolved with a gaussian kernel (640 samples, sd of 0.1) –Time-series y was generated as a product of weighted previous samples of x, according to:

$y_{t}$ = $\psi x_{t-n}$	(1),

where is the current value of y, is an autoregressive coefficient (set at 1 in this example) and is the previous sample of x. To keep the model simplistic, a model order of 1 was used, meaning that ‘y’ was generated using one previous sample of time series ‘x’. Both signals were then mixed with white noise with a standard deviation of 0.5. We simulated 100 trails in this way. In each trial, the amplitude of the sine wave used to generate the transient oscillation in x and y was set at 3 times a random number drawn from a normal distribution. This was to simulate the data being more physiologically ‘realistic’ as well as yielding an increase in the time domain and spectral GC, due to the amplitudes being highly correlated. Single trail time-frequency power was computed in the same way as in the simulation of concurrent power connectivity (see above). To compute GC, we used functionality from the MVGC toolbox (Barnett and Seth, 2014). In the accompanying code, we provide routines on implementing code from the toolbox with dual EEG data, to compute GC estimates for amplitude and power. As we generated the peak of signal y to be forward lagged by +100ms relative to the peak of x we can expect to see a relative increase in the time domain and spectral GC in a 100ms time window between peaks (in time) of signals x and y. We also expect that, as both x and y are transient signals and both signals return to baseline after 100ms, we will see no other significant GC increase. Further, as x was not defined by y, we expect a unidirectional influence only from x to y. The temporal precision of the GC results will be determined by the number of samples of the window used to compute GC. Previous literature has used window lengths between 50ms (Ding Bressler, Yang, and Liang, 2000) to 2s (Barret, Murphy, Bruno, Noirhomme, Bolfy, Laureys, and Seth, 2012) - although the most appropriate window length will depend on individual task-related design. In this example, GC was computed in a 200ms sliding window, in increments of 50ms. The model order used in the autoregressive model fit for the GC estimates was 1.

3. *Simulation for* ***non-event*** *locked changes in sequential amplitude/ power connectivity*

To illustrate how **non-event-locked** neural responses might give rise to changes in sequential amplitude/power IBC (see section 3.2.1 of main text) we simulated one time series as a mixture of one ‘slow’ sine wave at 2Hz and one ‘fast’ sine wave at 10Hz (640 time points, at a sampling rate of 256 Hz for both) and white noise (sd 0.5)- labelled as time series x. Time series y was generated as a product of weighted previous samples of x plus white noise with a standard deviation of 0.5, following the same procedure as in the previous simulation. In this example, a model order of 1 was used. We simulated 100 trials in this way**.** As y was generated from previous samples of x plus noise (random variation) to simulate a gradual increase in GC over time we linearly reduced the magnitude of noise parameter/ random variation in the system, such that the values of y become closer to the ‘true'/ real signal elements of x, without the confounding noise. GC was computed in the same way and using the same code as provided for the above simulation. The model order used in the autoregressive model fit for the GC estimates was 1.

4. *Simulation for event locked changes in concurrent phase connectivity*

To illustrate how event-locked neural responses might give rise to changes in concurrent phase-based IBC (see section 3.1.2 of main text), we simulated two-time series (640 samples at 256 Hz) as partially phase-locked signals (e.g. non-phase locked before time +200ms and purely phase locked after time = 200ms). At time=+200ms we simulated a phase-modulated such that at subsequent time points the signals were purely phase-locked. We simulated 100 trials of data in this way. To calculate the frequency-specific phase-locking value used in our examples (see section 3.1.2 of main text) both signals were filtered between 6 and 9 Hz using Matlab's window-based FIR filter with 625 points (equivalent to 3 times the lower edge of the bandpass range). The phase angles for both time series were taken from the result of the Hilbert transform. In the accompanying code, we provide routines on calculating inter-trial coherence (a measure of potential phase resetting) and phase locking between signals that can be applied to dual EEG data. In this example, PLV estimates were computed at each time points over trials (which gives the highest degree of temporal precision) though we also provide routines for calculating these same measures in a sliding window over time within trials.

5. *Simulation for event locked changes in sequential phase connectivity*

To illustrate how event-locked neural responses might give rise to changes in sequential phase-based IBC (see section 3.1.2 of main text) two-time series were generated as sine waves at 7Hz plus white noise (sd=0.5). For time series x, we simulated 100 trails of non-phase locked activity (e.g., by adding a random phase offset from the entire 0-2pi distribution). At time 0 we simulated a phase reset, whereby the phase was abruptly shifted. Following the phase reset (e.g., t=0 for signal x) the signals became phase-locked (e.g. with an added phase offset sampled from only half of the full 0-2pi distribution). Signal y was generated in the same way as signal x, but the 'event'/ phase reset was simulated 200ms after the phase reset in signal x. Phase angle again was obtained from the result of the Hilbert transform. Phase transfer entropy was computed in a 200ms sliding window

6. *Simulation for non-event locked changes in concurrent phase connectivity*

To illustrate how non-event-locked neural responses might give rise to changes in concurrent phase-based IBC (see section 3.2.2 of main text) we simulated two times series (640 samples at 256 Hz) x and y, as basic sine waves which varied in peak frequency over time. Time series x was designed to simulate an infant alpha generator with a frequency range of 6-9Hz and y an adult alpha generator with a frequency range of 9-12Hz. For the simulated infant data, the phase offset added at each time point to a 6Hz sine wave was designed as the cumulative sum of a series of spline interpolated, linearly spaced numbers *increasing* from 6 to 9 for the length of the time segment minus the mean of the upper and lower bounds of the frequency range. For the simulated adult data, the phase offset of the sine wave was set similarly, but a series of spline interpolated linearly spaced numbers *decreasing* from 12 to 9. We simulated 100 trials of data in this way. Phase angle again was obtained from the result of the Hilbert transform and PLV was calculated at each time point over trails as in the previous example.

7. *Time-frequency decomposition*

Single trail time-frequency power from the signals was derived by narrowband filtering the data between 2 and 25hz in section 2.3 of main text and 2 and 40Hz in section 3.1.1 in main text. 23/ 38 (respectively) filters were constructed, using MATLAB's FIR filter function, linearly spanning the frequency range. Each filter was constructed with an order of 3 times the sampling rate/ the lower limit of the frequency range (same as used by EEGLab), and with a frequency spread of +/- 1.5 Hz, giving a 2/3 overlap in frequencies between successive filters and then squaring the absolute values derived from the Hilbert transform of the filtered data. Time-frequency power and phase was then derived form the resulting Hilbert transformed data. For analysis involving time-frequency power, power was baseline normalised (decibel normalised) using activity in the -700 to -500ms time window and averaged over trials.

8. *Side note on inter trail phase coherence*

ITC measures the consistency of frequency band-specific phase angles over trials (time-locked to the response). The phase coherence value is computed according to:

$ITC=\left. \frac{1}{N} \right|\sum_{k=1}^{N} ⅇ^{i\phi\left( t,k \right)}$	(2),

where *N* is the number of trials, and is the phase angles of a signal on trial *n*, at time *t*, Phase coherence values vary from 0 to 1, where 0 indicates no phase consistency across trials to 1 indicates oscillations take on identical phase values across trials (Lachaux, Rodriguez, Martinerie, and Varela, 1999; Delorme and Makeig, 2004).

**References for supplementary materials**

Barnett, L., & Seth, A. K. (2014). The MVGC multivariate Granger causality toolbox: a new approach to Granger-causal inference. *Journal of neuroscience methods*, *223*, 50-68.

Barrett, A. B., Murphy, M., Bruno, M. A., Noirhomme, Q., Boly, M., Laureys, S., & Seth, A. K. (2012). Granger causality analysis of steady-state electroencephalographic signals during propofol-induced anaesthesia. *PloS one*, *7*(1), e29072.

Ding, M., Bressler, S. L., Yang, W., & Liang, H. (2000). Short-window spectral analysis of cortical event-related potentials by adaptive multivariate autoregressive modelling: data preprocessing, model validation, and variability assessment. *Biological Cybernetics*, *83*(1), 35-45.

Delorme, A., & Makeig, S. (2004). EEGLAB: an open-source toolbox for analysis of single-trial EEG dynamics including independent component analysis. *Journal of neuroscience methods*, *134*(1), 9-21.

Lachaux, J. P., Rodriguez, E., Martinerie, J., & Varela, F. J. (1999). Measuring phase synchrony in brain signals. *Human brain mapping*, *8*(4), 194-208.
